# Gene regulatory network that shaped the evolution of larval sensory organ in Cnidaria

**DOI:** 10.1101/2023.09.14.557727

**Authors:** Eleanor Gilbert, Jamie Craggs, Vengamanaidu Modepalli

## Abstract

Among non-bilaterian animals, a larval apical sensory organ with integrated neurons is only found in cnidarians. Within cnidarians, an apical organ with a ciliary tuft is mainly found in Actiniaria. Whether this apical tuft has evolved independently in Actiniaria or alternatively originated in the common ancestor of Cnidaria and Bilateria and was lost in specific groups is uncertain. We generated transcriptomes of the apical domain during the planula stage of four species representing three key groups of cnidarians: *Aurelia aurita* (Scyphozoa), *Nematostella vectensis* (Actiniaria), and *Acropora millepora* & *Acropora tenuis* (Scleractinia). We showed that the canonical genes implicated in patterning the apical domain of *Nematostella* are largely absent in *Aurelia*, indicating that scyphozoans and anthozoans do not share apical organ homology. In contrast, the apical domain of the scleractinian planula shares gene expression pattern with *Nematostella*. By comparing the larval single-cell transcriptomes, we revealed the apical organ cell type of Scleractinia and confirmed its homology to Actiniaria. However, *Fgfa2*, a vital regulator of the regionalisation of the *Nematostella* apical organ, is absent in the scleractinian genome. Likewise, we found that *FoxJ1* and 245 genes associated with cilia are exclusively expressed in the *Nematostella* apical domain, which is in line with the presence of ciliary apical tuft in Actiniaria and its absence in Scleractinia and Scyphozoa. Our findings suggest that the common ancestor of cnidarians and bilaterians lacked an apical organ with a ciliary tuft, and it could have evolved independently in several clades of cnidarians and bilaterians.

## Introduction

The majority of marine benthic invertebrates progress through a planktonic life phase during early development, a ciliated larva with an apical organ (1). The apical organ is a larval neurosecretory structure identified in the diverse groups of marine invertebrates and located at the frontal region of the larvae. Behavioural studies have demonstrated that ciliated larvae use the apical organ to process environmental cues and modulate their swimming behaviour (2-5). The apical domain of the larvae is often equipped with an ‘apical tuft’ of long cilia that protrude from apical cells (1, 6-12). Among non-bilaterians (Cnidaria, Placozoa, Porifera, and Ctenophora), a larval sensory organ with integrated neurons is only found in cnidarians **(Fig. 1A)** (13-15). The phylum Cnidaria (sea anemones, corals, hydroids and jellyfish) holds a key phylogenetic position for understanding the evolution of larval sensory-ciliomotor nervous system, providing an opportunity to investigate the primordial neurotransmission system.

**Figure 1:**
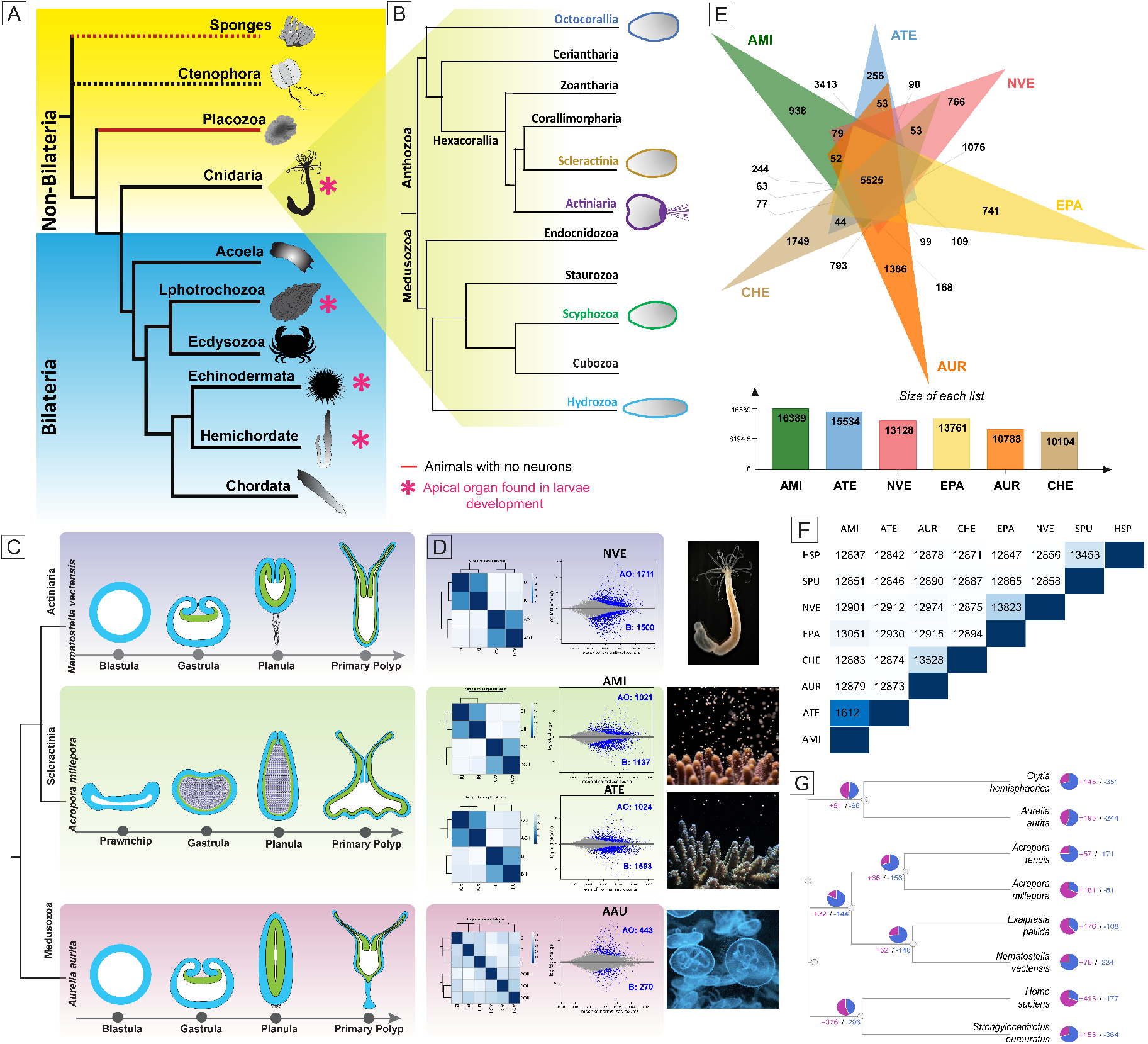
Evolutionary origin of the apical organ and spatial transcriptome of the cnidarian apical organ. (A) A brief overview of evolutionary relationships in the animal kingdom. Within non-bilaterians, a true larval apical organ with integrated neurons is only found in cnidarians. **(B)** An overview of evolutionary relationships in the phylum Cnidaria; an evident long ciliary apical tuft is commonly found in Actiniaria. **(C)** Schematic drawing of embryonic development in different cnidarian taxa (black–mesoglea, white–gastric cavity, green–endoderm, blue–ectoderm). The fertilised eggs progress through gastrulation and to planula. Planula larvae, after metamorphosis, transform into a feeding primary polyp. **(D)** MD and volcano plots represent the logFC ratio of differential expression between apical and body tissues from respective species. The significantly differentially expressed genes are highlighted in blue. MD plots display a global overview of all data sets. AO: apical organ; B: planula body. **(E)** A Venn diagram of orthologous genes shared between cnidarian species. The size of clusters in each species, including orthologues and in-paralogs. **(F)** Summary of proteins overlapped across each species. **(G)** The phylogenetic relationship of a selected list of cnidarians and bilaterian species was inferred based on orthologue gene groups and included the number of expanded and contracted orthologue groups indicated by CAFÉ analysis. Species abbreviations: AAU: *Aurelia aurita*, AMI: *Acropora millepora*, ATE: *Acropora tenuis*, CHE: *Clytia hemisphaerica*, EPA: *Exaiptasia pallida*, NVE: *Nematostella vectensis*, HSP: *Homo sapiens*, SPU: *Strongylocentrotus purpuratus*.

A more profound question is the evolutionary origin of apical organs, whether the apical organs of ciliated larvae across different phyla are homologous or evolved convergently. The morphology of the apical organ in cnidarian larvae is comparable to those of bilaterian larvae (2, 16, 17), indicating that the common ancestor of these two groups may have progressed through a free-swimming larval stage with a true larval apical organ and associated neurons (18, 19). A highly conserved set of genes paerning the apical/anterior ectoderm in Bilateria and in the apical/aboral ectoderm in cnidarian *Nematostella* (NVE) shows that these regions are very likely homologous (16, 20-24). This, in turn, makes it essenal to consider the possible “deep homology” of the apical organs across different phyla. Given the remarkable fracon of transcriptional factors (TFs) of the apical gene regulatory network (GRN) that also contribute to the development of other nervous systems of various forms and funcons (25-29), it is conceivable that regulatory modules that are commonly expressed in animals with more complex nervous systems are also deployed in cnidarian larvae. Strikingly, many TFs associated with nervous system development have demonstrated a conserved spaal distribuon across the anterior-posterior axis of the nervous system among cnidarian NVE and bilaterian species (27, 28). Whether these homologous TFs can also be recognised across different cnidarian taxa is undefined.

Cnidarians are divided into two major groups: Anthozoa (sea anemones, corals and sea pens) and Medusozoa (jellyfish, sea wasps and Hydra) **(Fig. 1B)**. Even though cnidarians are monophyletic, the magnitude of genetic differences between the anthozoans and medusozoans is equivalent to that between Anthozoa and Deuterostomia (30, 31). Molecular dating estimated the separation of the major cnidarian clades more than 500 MYA, and each group has undergone a long period of independent evolution (31). Though a large proportion of early embryonic development is shared across these groups, the actiniarian planula, like NVE, is morphologically unique as it possesses a long ciliated apical tuft **(Fig. 1B)**. The apical domain displays several apical cells bearing a ciliated tuft as well as RPamide and PRGamide peptidergic cells (32, 33). In Medusozoa and Scleractinia, the apical domain is also enriched with neuropeptide-expressing cells (3, 34-37). However, unlike Actiniaria, the Scleractinia and Medusozoa commonly lack a long ciliated tuft. Exceptionally, some non-reef building coral species in the Rhizangiidae family of Scleractinia have been documented with ciliary apical tuft, although this has not been published. The widespread occurrence of apical organ in multiple phyletic groups of marine invertebrates **(Fig. 1A, B)** and clade-specific absence has prompted radically different views on the origin of the apical organ. Some consider them an ancient feature of the eumetazoan ciliated larvae (38-41). Alternatively, others argue that they could have evolved multiple times independently (42-45). Within cnidarians, the inconsistency of apical organ structures and the lack of apical tuft in the frontal region of Scleractinia and jellyfish larvae has raised doubt over the common origin of the apical organ and the advantages of long ciliated tuft in specific groups. **(Fig. 1B)**. Whether the apical organ with a tuft is ancestral or secondarily evolved and which GRN facilitated the evolution of the apical tuft is yet to be addressed.

Here, we employed a spatial transcriptomic approach to compare the molecular basis of the apical domain in three groups of cnidarians, including Actiniaria (sea anemones), Scleractinia (stony corals) and Scyphozoa (true jellyfishes). This technique has proven robust in identifying the apical enriched genes in our previous study in NVE (33). We examined the spatial distribution of transcription factors and other key signalling components previously studied in the frontal/apical region of the larvae (2, 33, 46-53). We revealed the evolutionary relationship of apical organ between Cnidaria groups.

## Results and discussion

### Tissue-specific transcriptome of cnidarian planula reveals the molecular topology tightly associated with the apical domain

To investigate the molecular topology of the apical domain among cnidarian groups, we systematically carried out microdissections on cnidarian larvae collected at the planula stage from Actiniaria *Nematostella vectensis* (NVE), Scyphozoa *Aurelia aurita* (AAU) and two Scleractinia species *Acropora millepora* (AMI) and *Acropora tenuis* (ATE) (**Fig. 1C**). We carefully separated the apical tissue from the rest of the larval body and acquired transcriptomic data from both the apical and the rest of the body tissues separately to perform differential gene expression (DGE) analysis. By using DGE analysis, we idenfied significantly differenally expressed gene (DEGs) between the apical and body ssue of each species. **(Fig. 1D)**. The global gene expression patterns among the apical and body ssues from replicates were compared using principal component analysis and correlation analysis (**Fig. 1D**); the plots displayed a strong correlation among the replicates. Notably, *Aurelia* presented a relatively low number of significantly DEGs: 713 (padj (FDR) < 0.05). Of all four species, the Actiniaria NVE presented the highest number of significantly DEGs (3225) between apical and body tissues. The number of significantly DEGs for each species is presented in **Figure 1D, Supplementary Table 1**.

### Orthology analyses to compare the planula anteroposterior molecular composition between Actiniaria, Scyphozoa and Scleractinia larvae

To enable comparison of planula anteroposterior patterning among different groups of Cnidaria, we reveal the molecular composition of anteroposterior domains in NVE, AMI, ATE and AAU using the spatial gene expression data. First, by using orthology analyses, we grouped the cnidarian proteomes to identify both homologous and species-specific genes **(Fig. 1E-G)**. Along with NVE, AMI, ATE and AAU, we additionally included cnidarians *Clytia hemisphaerica* and *Exaiptasia pallida* as well as bilaterians *Strongylocentrotus purpuratus* and *Homo sapiens*. The output data from the orthology analyses are provided in **Supplementary file 1**. Next, we sorted the genes enriched in apical and body tissues from DEGs data into two groups. Using OrthoVenn3, we identified orthologous groups shared across four species in apical and body tissues separately **(Fig. 2)**. Within apical tissue enriched gene sets, 58 orthogroups with 314 proteins were expressed in all four species, 152 orthogroups (522 proteins) were shared across anthozoan species, and 261 orthogroups (540 proteins) were shared exclusively between Scleractinia AMI and ATE. 92 orthogroups (206 proteins) were expressed only in the NVE **(Fig. 2B)**. NVE also expressed 1027 singletons exclusively in the apical domain. As detailed in Figure 2C, we also defined co-expression dynamics of genes in the posterior domain or body tissue across NVE, AMI, ATE and AAU through orthology analyses **(Fig. 2C)**.

**Figure 2:**
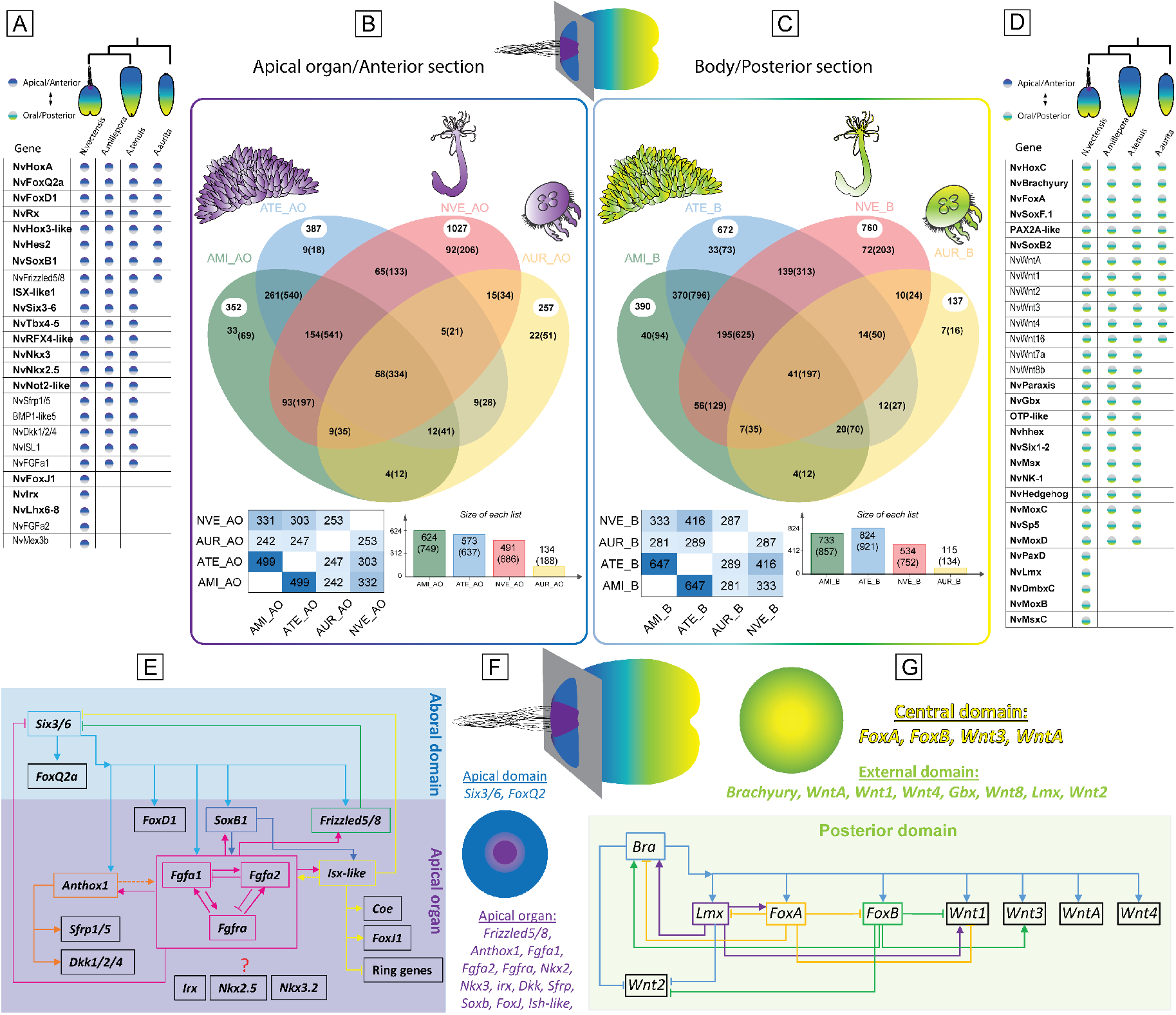
Comparison of tissue-specific genes with shared expressed between cnidarian planulae. (A, D) The table detailing each TF (bold) and other critical developmental genes enriched in apical domain **(A)** and body section of planula **(D)** from NVE, AMI, ATE and AAU. **(B, C)** Venn diagrams presented the distribution of shared and unique orthogroups between NVE, AMI, ATE and AAU in apical domain (B) and the body section of planula (C). The number of proteins in ortholog groups/clusters are indicated in brackets next to the number of ortholog groups/clusters. The number in white circles indicates the number of singletons. At the boom presented the summary of proteins overlapped across each species and the size of clusters in each species, including orthologues and in-paralogs. **(E-G)** Illustrang NVE apical domain and oral/posterior domain GRN (2, 15, 20, 21, 33, 47, 52, 54-57). Segregaon of expression domains in NVE planula apical domain divided into ring (blue) and spot (purple) territories (2, 21, 33, 54, 57). Likewise, in the posterior region, a set of genes are enriched in the central domain (yellow), and others adjust to oral (green) (15, 20, 58).

### Despite the lack of apical tuft, scleractinian planulae share conserved anteroposterior patterning genes with Actiniaria

Scleractinian corals belong to the Hexacorallia, a lineage within the class Anthozoa and a sister group to the Actiniaria **(Fig. 1B)**. Yet, the scleractinian frontal region differs from the anthozoan planula mainly the ciliary apical tuft is absent. The extent to which the Scleractinia apical domain shares molecular topography with its sister group Anthozoa is yet to be addressed. The frontal region of cnidarian embryos is set by an apical domain GRN directing the formation of apical organ. The well-defined developmental GRN in NVE provides a powerful framework for investigating the evolution of embryonic patterning mechanisms (14, 20, 33, 46, 49, 52, 53, 56, 57, 59). If the apical domain is evolutionarily related across cnidarians, an overlap of GRNs would be expected. Using spatial transcription data and comparative genomics, we first sought to define the co-expression dynamics of genes involved in the patterning anteroposterior territories along the planula apical and body tissues **(Supplementary Table 1)**.

In NVE, the apical domain GRN is composed of *NvIsx-like, NvFgfa1, NvFgfa2, NvSfrp, NvFrizzled-like, NvDickkopf-like, NvNkx3, NvNkx2*.*5, NvTbx4-5, NvHoxA* (*Anthox6*), *NvFoxQ2a, NvIrx, NvFoxD1, NvRX, NvHox3-like, NvHes2, NvSoxB1, NvSix3-6, NvRFX4-like, NvFoxJ1, NvTauD, PoxA-like* and *Bmp1-like5* genes. Chiefly, *NvHoxA* (*Anthox6*), *NvFgfa1, NvFgfa2, NvSix3-6*, and *NvIsx-like* have been functionally studied, and the knockdown (KD) of each of these genes independently has resulted in the ether loss of apical tuft cells or defects in apical organ development (33, 49, 54). Using spatial transcription data, we compared the shared expression of these orthologues in the Scyphozoa and Scleractinia planula apical domain. As illustrated in **Figure 2A**, genes that pattern apical domain of NVE are largely expressed in similar patterns in Scleractinia AMI and ATE planula this includes *Isx-like, Fgfa1, Fgfa2*,, *Sfrp, Frizzled-like, Dickkopf-like, Nkx3, Nkx2*.*5, Tbx4-5, HoxA* (*Anthox6*), *FoxQ2a, FoxD1, Rx, Six3-6, Rfx4-like, FoxJ1, Hes2, SoxB1, PoxA-like* and *Bmp1-like5* genes. Despite the lack of apical tuft, Scleractinia planulae share a similar set of genes in their apical domain, suggesting that these TFs already operated as a part of a GRN that was assembled before the separation of Actiniaria and Scleractinia.

In parallel, we also compared the genes associated with posterior territory using NVE GRN as a template **(Fig. 2)**. We compared the posterior region with the genes including *NvHoxC, NvBarH, NvFoxA, NvSoxF, NvGbx, NvMsx, NvHedgehod, NvMoxA, NvMoxB, NvMoxC, NvMoxD, NvPaxD, NvLMX*, and members of the *Wnt* (*NvWntA, NvWnt1, NvWnt3, NvWnt4, NvWnt16, NvWnt2* and *NvWnt8*). Note that with some of the TFs, we could only find significant differential expression in one of the *Acropora* species; therefore, we excluded them from the list **(Supplementary Table 1)**. As illustrated in **Figure 2D**, genes that pattern the posterior territory of NVE are largely expressed in similar patterns in Scleractinia AMI and ATE planula, including *HoxC, BarH, FoxA, SoxF, Gbx, Msx, Hedgehog, MoxC, MoxD*, and members of the *Wnt* family (*WntA, Wnt1, Wnt3, Wnt4, Wnt16, Wnt2* and *Wnt8*). This reveals that Scleractinia and Actiniaria share a large portion of GRN in both the anterior and posterior domains.

The cellular identity of apical organ has never been detailed in Scleractinian larvae. A previously published study of larval single-cell transcriptome in the scleractinian *Stylophora pistillata* has not classified apical organ cell types (60). From the spatial transcriptome and orthology analyses, we revealed the shared apical domain genes in scleractinian planula. Using characterised NVE apical organ marker genes and single-cell transcriptome data (61), in combination with orthology analyses, we investigated the Scleractinia planula single-cell data (60) to predict apical cell types. We first identified the orthologues shared across NVE, AMI and *Stylophora pistillata* **(Fig. 3A)**. Next, based on previous studies in NVE, we selected apical organ genes including *NvIsx-like, NvFgfa1, NvFgfa2, NvFgfra, NvSfrp, NvTauD, NvFrizzled-like, NvDickkopf-like, PoxA-like, Bmp1-like5, NvNkx3, NvNkx2*.*5, NvLhx6-8, NvTbx4-5, NvHoxA* (*Anthox6*), *NvFoxQ2a* and *NvIrx* **(Fig. 3B, E)**. We also included genes expressed in the apical domain enriched cell types such as gland cells, larval specific neurons and undifferentiated cell type 3: *NvRx, Hmcn1-like5, NvSix3-6, Klkb1-like3, NvRfx4-like, Anxa6-like, Nvcreb-like2, Dbp-like2* and *NvTrerf1-like* **(Fig. 3C)**. Next, we pulled out the expression patterns of the list of genes associated with the NVE apical organ. From the comparison, we identified a set of apical organ genes that are specifically expressed in an undefined cell type 22 **(Fig. 3D)**. This cell type displayed shared expression of some of well characterised NVE apical organ genes, including *Frizzled-like, Sfrp, Lrp5/6, Isx-like, Dickkopf-like, Nkx3, TauD, Fgfa1, Fgfra, Bmp1-like5, Tbx4-5, Spon1-like*, and *PoxA-like* **(Fig. 3D)**. This analysis reveals the apical cell types (cluster 22) in *Stylophora pistillata* and provides additional evidence on shared apical organ cell types between actiniarian and scleractinian species.

**Figure 3:**
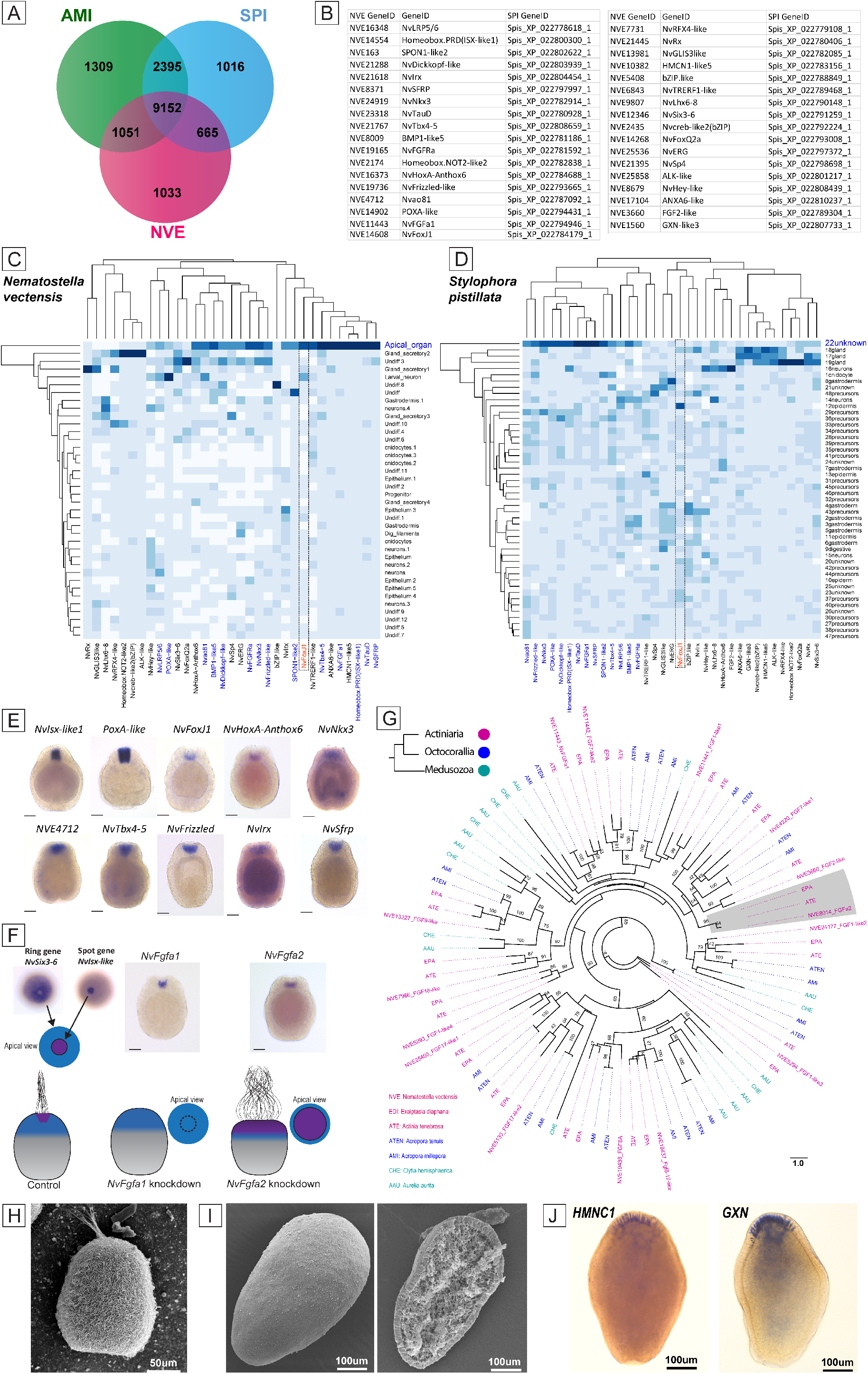
Defining apical organ in Scleractinia planula. (A) Venn diagrams presenting the distribution of shared and unique orthogroups between NVE, AMI and SPI. **(B)** A table detailing the homologs of NVE apical domain genes identified in *Stylophora pistillata* and their respective gene IDs. **(C, D)** Heatmaps displaying the gene expression of selected apical domain marker genes across larval cell types classified through single-cell transcriptomes of NVE (C) and *Stylophora pistillata* (D). **(E)** ISH of apical organ-enriched TFs. **(F)** Schematic drawings illustrating the morphological phenotype after *NvFgfa1* and *NvFgfa2* KD. *NvFgfa1* KD produces larvae lacking an apical organ and ciliary tuft, while *NvFgfa2* KD leads to larvae with an expanded apical organ and ciliary tuft. **(G)** A phylogenetic relationship of cnidarian FGF proteins. Values on nodes represent Bootstrap values (100 replicates). Bootstrap support values above 50% are indicated above branches. For additional details, refer to **Supplementary Table 1. (H)** SEM of NVE planula viewed laterally. **(I)** SEM of AMI planula viewed laterally, images on the right; the planula was cracked open to visualise the internal structures. **(J)** The expression of the top two genes *HMNC1* (XP_044178031.1) and *GXN* (XP_029194558.2), enriched in the AMI apical domain, was determined by ISH.

Of all the apical domain genes, we noted that *Fgfa2*, a critical regulator of NVE apical organ, is primarily absent in the AMI and ATE differential expression data. Further, from the FGF phylogenetic analysis, we identified that *Fgfa2* is absent in the Scleractinian genomes **(Fig. 3G)**. The development of the apical ciliary organ in NVE is under the control of two paralogous FGF genes (*NvFgfa1* and *NvFgfa2*), and one FGF receptor gene (*NvFgfra*) are expressed at the apical domain of NVE with spot expression pattern (21, 49) **(Fig. 3F)**. As illustrated in **Figure. 2E**, the aboral domain surrounding and encompassing the apical tuft structure reveals two major concentric domains. The first external circle in the apical domain expressing *NvSix3/6, NvFoxQ2a*, and *NvFoxD1* are devoid from apical tuft/apical organ territory **(Fig. 2E, blue colour)**. While *NvFgfa1, NvFgfa2, NvFGFRa, NvIrx, NvFoxJ* and *NvIsx-like* are restricted mainly to the spot region of apical domain, this also includes other spot genes *Dhh, NvSfrp1/5, NvNkx2*.*1, NvNkx3*.*5, NvSoxB* and *NvHoxF/Anthox1* (21) **(Fig. 2E, purple colour)**. KD experiments show that the signalling of *NvFGFa1* is required for the specification of the apical organ’s ciliary tuft. In contrast, *NvFgfa2* is necessary to limit the size of ciliary tuft cells to spot regions by the antagonistic interplay of *NvFgfa1* and *NvFgfa2* signalling **(Fig. 3F)** (21, 49). The morphology of the scleractinian planula comparatively differ from at the frontal region from that of the anthozoan planula and somewhat resembles a *NvFgfa2* KD planula **(Fig. 3F, H-J)**. Similarly, based on the expression of the top two genes HMNC1 and GXN, enriched in the AMI apical domain, the cells in this region displayed a wide distribution. It seems, *NvFgfa2* is an actiniarian innovation to restrain the apical cells to spot area of the apical domain and this extensive patterning of the apical domain is unique to anthozoan clade. However, a Further analysis of the spatial distribution of apical cells in Scleractinia planula is necessary.

### GRN that shaped the actiniarian apical organ with long ciliated tuft

*FoxJ1* is the master regulator of ciliogenesis (62-64) and orthologues of *FoxJ1* have been found in vertebrates and invertebrates. TFs, such as FGF (59) and Wnt (65), act as upstream regulators of *FoxJ1* ciliogenesis (16). A conserved role of *FoxJ1* in motile cilia formation is supported by expression patterns outside chordates. In sea urchin (phylum Echinodermata) larvae, *FoxJ1* expression has been shown in the most apical ectoderm marking the apical tuft (66-68). Similarly, in annelid *Platynereis dumerilii*, the *FoxJ1* is expressed in the apical plate and the ciliated bands. In NVE, along with other spot genes, *NvFoxJ1* expression is observed in spot region **(Fig. 3E)** and its expression coincident with the development of the apical tuft (69). Like in bilaterians, the *NvFoxJ1* expression in NVE is under the control of FGF singling **(Fig. 2E)**. KD of *Fgfa1* affects the expression of *NvFoxJ1*, which affects the apical tuft cilia (49). Unlike NVE, the apical domain of Scleractinia and AAU planula lack enrichment of *FoxJ1* expression. Further, using the larval single-cell data, we compared *FoxJ1* expression among scleractinian *Stylophora pistillata* (60) and actiniarian NVE larval cell types (61). As indicated in **Figure 3C, E**, in NVE, the *NvFoxJ1* is significantly expressed in the apical organ cell type. In contrast, in *Stylophora pistillata* the *FoxJ1* is enriched in the epidermis but not in the predicted apical cell type (cluster 22 unknown) **(Fig. 3D)**. Jointly, these findings suggest that *FoxJ1* expression apical organ of anthozoans is unique and likely supporting apical tuft formation. However, functional studies will have to determine whether *NvFoxJ1* is specifically required for the development of the apical tuft cilia or whether the motile cilia of the body surface also depend on the function of this gene.

Finally, we investigated if Actiniaria exhibits unique expression of ciliary genes in the apical domain compared to scleractinian and medusozoan planula. The apical tuft cilia are found in a wide range of marine invertebrates such as echinoderms (70), molluscs (71), annelids (72, 73) and cnidarians (33). In addition to the ciliary tuft, the ciliated larvae exhibit motile cilia. In general, the motile cilia typically cover the entire body of the larvae, aiding in the organism’s movement through ciliary beating **(Fig. 4A)** (1). The apical tuft does not contribute to larval motility, however, the ciliary tuft in NVE displays movement by expanding and contracting into a ciliary bundle. A comparative study of the ciliary proteomes in NVE planulae, sea urchins, and choanoflagellates (94) revealed core components of the ciliary intercellular signalling pathways and identified the shared ciliary proteome. The ciliary proteome data were acquired by isolating cilia from whole NVE planula, which were subjected to mass spectrometry; this allowed the construction of the ciliary proteome from the whole larvae, including the apical tuft (74).

**Figure. 4:**
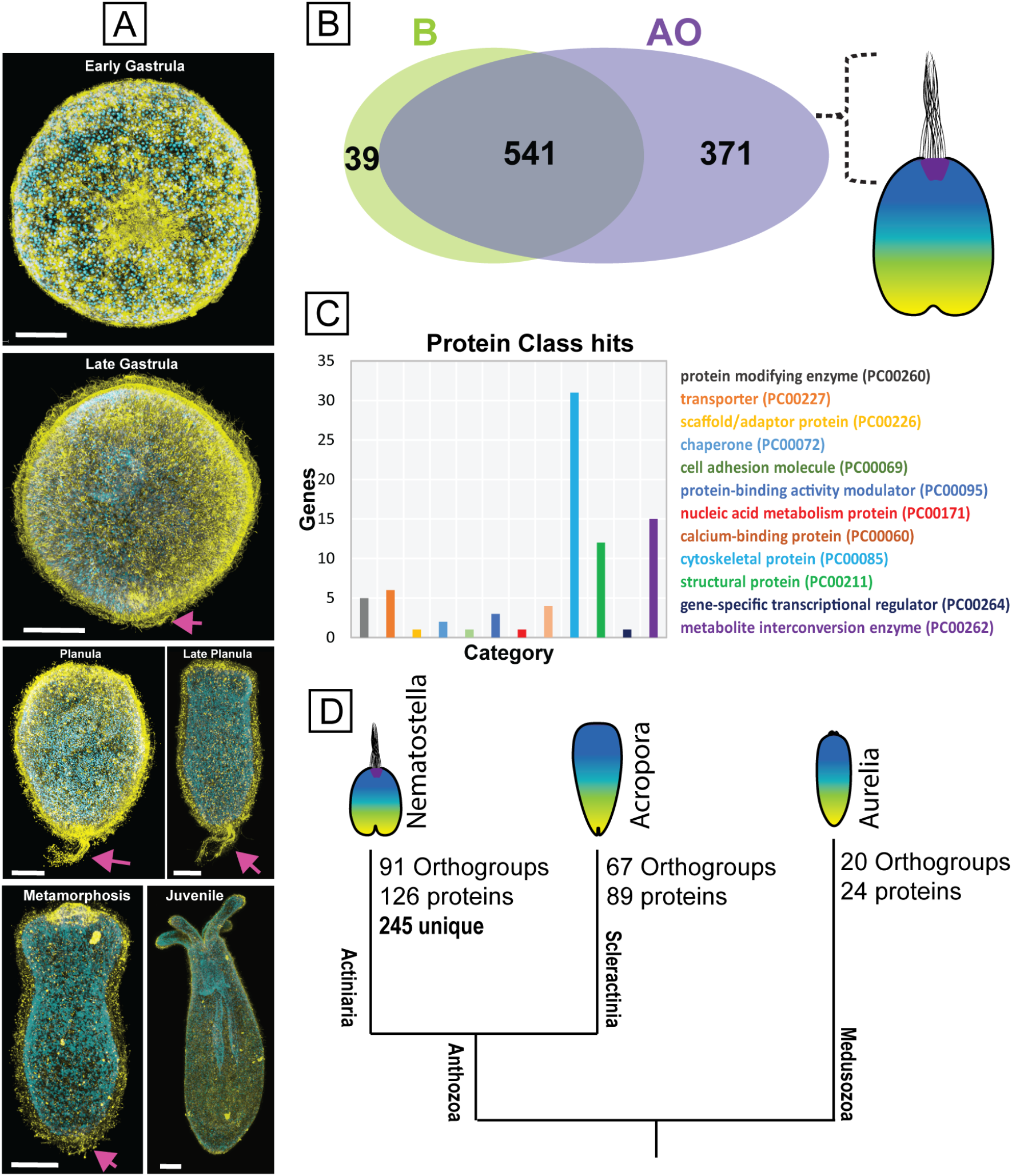
The genes linked with ciliary tuft in Cnidaria. (A) Different stages of *Nematostella* development. Immunostaining (yellow) with the acetylated tubulin antibody and counterstaining with DAPI (blue) for nuclei. An arrow pointing to apical tuft. **(B&C)** Tissue-specific transcriptome vs ciliary proteome of NVE planula to define cilium-associated genes enriched in apical tuft. **(B)** Venn diagram showing the DEG from the body and apical tissues. **(C)** The apical enriched cilia genes are categorised into different protein classes using gene ontology. **(D)** NVE apically enriched cilia orthologues shared with AMI, ATE and AAU. Out of 371 apically enriched cilia genes, NVE uniquely expresses 245 cilia-specific genes in apical domain.

Hear, we integrated the tissue-specific transcriptomes with cilia proteomes from NVE planula **(Fig 4B) Supplementary Table 1**. Among the DE genes, 371 ciliary genes were enriched in the apical region, 541 ciliary genes were commonly expressed throughout the body, and 39 were significantly enriched in the body, suggesting that the apical tuft cilia possess a set of proteins distinct from the rest of the body cilia. Among the apical enriched genes, we came across a set of candidates related to cilium organisation, cytoskeletal and structural proteins **(Fig. 4C)**, such as dynein heavy chain, axonemal, beta-tubulin, bardet-biedl syndrome, kinesin family member, enkur, tektin ADP ribosylation factor like GTPase, filamin A, tetratricopeptide repeat domain, stabiliser of axonemal microtubules 1, usherin, and intraflagellar transport protein. We also observed genes associated with metabolite interconversion such as WD repeat-containing protein and kinase family members (phosphoenolpyruvate carboxy kinase, nucleoside diphosphate kinase, adenylate kinase) **Supplementary Table. 1**.

Lastly, to investigated if Actiniaria exhibits unique expression of ciliary genes in the apical domain compared to scleractinian and medusozoan planula. Using orthology analysis and spatial transcriptome data we compare the NVE apical enriched ciliary genes among AMI, ATE and AAU apical tissue data **(Fig. 4D)**. Out of 371 ciliary genes enriched in the apical domain of NVE, 245 genes were undetected in apical domain of AMI, ATE and AAU apical tissue **Supplementary Table 1 and File. 1**. This suggests that Actiniaria planula innovated a large set of ciliary genes leading to the origin of a unique apical organ with a ciliary tuft. However, it should be noted that some of those 245 ciliary gene orthologues may present in the genomes of Scleractinia and Medusozoa, but not expressed specifically in apical domain of planula stage. The crucial factor is the spatial-temporal expression of cilia genes, specifically in the apical domain during the planula stage, which is essential for the development of apical tuft cilia in the Actiniaria.

### A distinct molecular topology defines the *Aurelia* apical domain: most of the canonical TFs associated with apical organ GRN are absent in AAU

The Medusozoa is a sister group to the Anthozoa **(Fig. 1B)**. To address the nature of the molecular differences between NVE and AAU, we investigated ortholog data to identify the shared apical domain genes. We found significant expression of *Hox9-14C, FoxD1, Rx, SoxB1, Hes2* and *Frizzled* regulatory genes in the AAU planula apical domain, while *Isx-like, Fgfa1, Fgfa2, Irx, Six3-6, Nkx3, Nkx2*.*5, Tbx4-5, Sfrp, TauD, Dickkopf-like, Bmp1-like5, Isl1, Erg1* and *Mex3b* genes had no significant differential expression **(Fig. 2D)**. Genes like *Fgfa1* and *Six3-6* showed a lack of minimal read count. Through phylogenetic analysis we confirmed that both *Fgfa2* and *Isx-like* are absent in the genome of Medusozoa **(Fig. 3E, 4A)**. At least from the orthology analyses, we did not find the orthologues of *Nkx2*.*5, Not2-like, Rfx4-like* and *Bmp1-like5* in the genome of AAU. Here we discuss the criticality of some of these genes to highlight their key role in the apical domain and the evolutionary consequences for the apical organ.

*NvSix3/6* has a broad role in determining the identity of the aboral domain development. The *Six3/6* antagonistic role in repressing *Wnt* singling allows the activation of aboral genes, including *FoxQ2, Rx*, and *Nkx* (75). KD of *NvSix3/6* affects the expression of several apical domain genes, including *NvFGFa1, NvFrizzled5/8* and *NvFoxQ2a* (21), which are in turn associated with the expression of downstream genes *NvIrx, NvIsx-like, NvFgfa2* and *NvFoxJ1* genes, together resulting in a loss of the apical organ and further affecting the larval development **(Fig. 2E)** (21, 47). Likewise, in bilaterians like sea urchins, KD of *six3* function results in a loss of the apical plate and impairs neural development (75, 76). Strikingly, in AAU, along with *six3/6*, many of its downstream genes, including *Irx, Isx-like, Nkx3, Nkx2*.*5, Dkk, Fgfa1, Fgfa2* and *FoxJ1* genes either lack shared spatiotemporal expression with NVE apical domain or absent from the genome of Medusozoa **(Fig. 2A, 4B)**.

Previously in NVE, we identified *NvIsx-like*, a PRD class homeobox gene expressed explicitly in apical tuft cells, as an FGF signalling-dependent TF responsible for forming the apical organ (33). *Isx-like* KD prevented the formation of the apical tuft cilia and loss of the apical tuft cell identity (33). From the phylogenetic analysis of the PRD class homeobox gene, we identified that *NvIsx-like* is absent in the medusozoans genome **(Fig. 5A)**.

**Figure 5:**
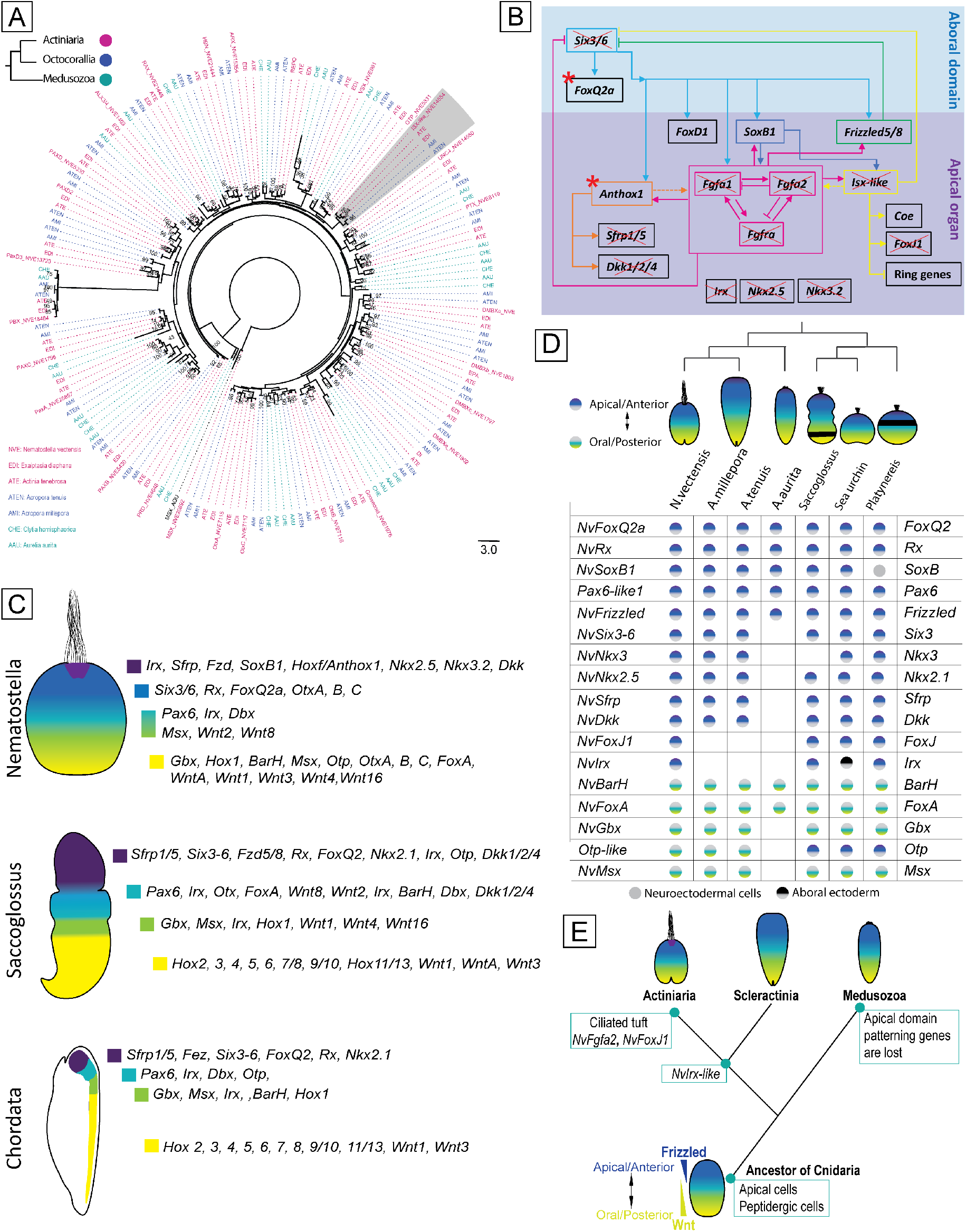
A distinct molecular topology of *Aurelia* apical domain and lack of shared expression of putative neural patterning genes. (A) A phylogenetic relationship of cnidarian PRD class proteins. Values on nodes represent Bootstrap values (100 replicates). Bootstrap support values above 50% are indicated above branches. **(B)** Illustrating the apical domain GRN of NVE and superimposing the AAU apical domain genes scenario. The genes that are absent in the apical domain are indicated with a cross. *FoxQ2* and *Anthox1* are expressed but not homologous to NVE. **(C)** Expression of nervous system axial patterning genes across cnidarians and bilaterians (16, 21, 27, 28, 67, 77-83). Schematic representation of the anteroposterior expression domains of genes in cnidarian NVE and a similar set of genes known to deploy in anteroposterior neuronal patterning in bilaterian species. **(D)** Shared expression of anteroposterior patterning genes in the larva of cnidarian and bilaterian invertebrate species. Most genes conserved across cnidarian NVE, AMI, ATE and bilaterian species lack shared expression in AAU. **(E)** A possible scenario of apical tuft origin in Cnidaria: the common ancestor of cnidarians possessed apical plates comprised of apical and sensory neurosecretory cells. Apical cells homologues to anthozoans and bilaterians may have lost in the Medusozoa after splitting from Anthozoans. After the split of Actiniaria from other Anthozoans, the Actiniaria innovated genes like *NvFgfa2* and a set of ciliary genes that added a long ciliary tuft to the apical organ.

Unlike *Isx-like* homeobox gene, AAU showed another *Hox* gene expression in the apical domain **(Fig. 2A)**. From the orthology analysis we initially noticed it clustering with NvAnthox1 orthogroup. However, from previous phylogenetic studies (84, 85), we understand that it is actually paralogous to NvAnthox1. NvAnthox1 is notable for its apical domain expression in NVE (86). A comparative study across anthozoans and medusozoans phylogenetically placed the anthozoan NvAnthox1 clade as a sister to a pair of medusozoan Hox clades, including Hox9-14C (84, 85). Previous studies in hydrozoan species, including *Clytia hemisphaerica* and *Cassiopea xamachana*, have shown *Hox9-14C* expression in apical domain (84, 85). Similarly, the spatial transcriptome of AAU apical domain showed significant expression of a *Hox* gene homologous to *Hox9-14C*. The consistent pattern of aborally localised expression of *Anthox1/Hox9-14C* in both anthozoans and medusozoans suggests that the cnidarian ancestor utilised *Anthox1/Hox9-14C* signalling in apical domain GRN.

Another well-conserved marker for apical domain territories is the forkhead domain transcription factor *FoxQ2* **(Fig. 3E)**, which functions downstream of *Six3/6* in the development of the apical domain of NVE **(Fig. 2E)** (21) and sea urchins (75). In AAU, unlike *Six3/6, FoxQ2* shows enrichment in the apical domain. However, as previously identified in *Clytia*, the AAU apical enriched *FoxQ2* gene is not an orthologue of NVE *NvFoxQ2a* (87, 88). The GRN around medusozoan *FoxQ2* is yet to be studied. Overall, most of the canonical GRN associated with apical organ lack significant expression or is missing from the AAU genome. Some of these findings also coincide with previous studies in the hydrozoan species *Clytia hemisphaerica* (85, 87, 88), belonging to a separate subgroup of Medusozoa.

### A conserved neuronal expression domain map between cnidarians Anthozoan and bilaterians, but not in Medusozoa AAU

The frontal region of cnidarian larvae is set by an apical domain GRN directing the formation of apical organ and subsequent specification of neurons. Developing nervous systems are regionalised by stripes of gene expression along the anteroposterior axis. Along with apical nervous system, the NVE planula simultaneously develops neurons at the blastopore/oral end (13, 28). Numerous TFs show concentric expression in the oral ectoderm of NVE planula larvae, anticipating in development of oral/blastoporal neurons (14, 47, 52, 89) **(Fig. 5C)**. Notably, many of these TFs are well-established players in patterning of bilaterian neuroectoderm (90-92), expressed in a similar sequence of domains in annelids (91), cephalochordates (29, 93) and hemichordates (82). The expression of nervous system genes can be segregated into three broad groups to facilitate the comparison between cnidarians and bilaterians: anterior, midlevel, and posterior genes. As diagrammed in the **Figure 5C**, at the anterior domain, 13 genes were identified in NVE, namely *NvSix3/6, NvFoxQ2, NvRx, NvFoxD1, NvOtx, NvFrizzled5/8, NvNkx2*.*5, NvNkx3*.*2, NvIrx, NvDkk, NvSfrp1/5, NvSoxB1* and *NvFoxJ1*. Most of these genes are known to express within the forebrain or anterior territory of hemichordates (26) and they all have prominent expression domains in the prosome ectoderm of *S. kowalevskii*, the hemichordate’s most anterior body part **(Fig. 5C)** (80, 94, 95); these ortholog cognates express entirely within the apical domain in NVE. Midlevel genes are those expressed in the mesosome and anterior metasome (with some domains extending anteriorly into the prosome), that is, more posteriorly than those genes of the anterior group. In case of hemichordates, they expressed at least in the midbrain, having posterior boundaries in the midbrain or anterior hindbrain **(Fig. 5C)**. In NVE, at the midlevel ten genes were identified, namely *NvPax6, NvIrx, NvDbx, Nvlim1/5, NvMsx, NvDlx, NvDll, NvWnt2* and *NvWnt8*. Some of these genes differ in their anterior and posterior extent **(Fig. 5C)**. Posterior genes are those expressed entirely within the hindbrain and spinal cord regions of the chordate nervous system. In hemichordates, they share orthologues expression in the posterior metasome (79, 80, 94). At the posterior domain, 12 genes were compared, namely *NvGbx, NvBrah, <u>NvMsx</u>, NvOtp, NvOtx, NvFoxA, NvHox1, NvWntA, NvWnt1, NvWnt3, NvWnt4* and *NvWnt16* as shown in the **figure. 5C**. All these genes are expressed in the oral region of NVE. Despite the significant phylogenetic distance, the relative order in which the axial TFs are expressed in NVE is similar to the order of their expression along the axis of the bilaterian nervous system development **(Fig. 5C)**.

Taking these nested concentric domains of NVE genes, we asked whether these orthologous genes are expressed in a similar pattern along the anterior-posterior axis in other cnidarian groups (96, 97). Using spatial transcriptome, we can only distinguish the expression of a gene enriched in the apical domain or the rest of the body. Therefore, to facilitate the comparison, we combined the midlevel and posterior genes in AAU, AMI and ATE. As shown in the **figure. 5D**, we indicated the list of shared neuronal genes across the anterior and midlevel/posterior domains of planula. Comparing the orthologues between Actiniaria and Scleractinia larvae show majority of genes with in similar spatial expression pattern **(Fig. 5D)**. In contrast, the AAU larvae lack shared expression of *Irx, Isx-like, Six3/6, Nkx3, Nkx2*.*5* and *Dkk* genes, while *FoxQ2, Rx, Frizzled5/8, Sfrp1/5, SoxB1, Brah, FoxA, Wnt1, Wnt3, Wnt4* and *Wnt16* have shown the similar expression pattern with NVE **(Fig. 2D, 5D)**. Thus, in bilaterians, as in cnidarians, *Gbx, Msx, BarH, Pax, Dbx, FoxA, FoxB* and *Lim* members are expressed in the lateral region. In the case of AAU, the *Msx* and *Dbx* are absent in posterior domain, whereas *Gbx* and *Pax3-7/PaxD* are absent in the genome (31, 98). Along both anterior and posterior domains, the AAU showed variations in GRN; several crucial apical organ and neuronal genes along the anteroposterior axis are lacking conserved expression with NVE. Hence, most of these genes are evolutionarily conserved between anthozoans and bilaterians **(Fig. 5C)** and are associated with anteroposterior neurogenesis and axial formation suggesting that AAU has undergone extensive GRN changes after the anthozoans split. Therefore, our findings demonstrate that the magnitude of molecular differences between medusozoans and anthozoans is greater than those between anthozoans and bilaterian invertebrates. **(Fig. 5D)**.

## Conclusion

The inconsistency of the apical organ with ciliary tuft among cnidarians provides a window to understand the evolution of apical organ and the GRN that operated in the common ancestor of eumetazoans. Comparative gene expression studies between the cnidarian NVE and bilaterian ciliated larvae revealed a strong resemblance in the molecular topography around the apical pole (16, 21, 28, 99), suggesting that the apical organ may be an evolutionarily conserved larval structure and might have appeared within the ancestor of Eumetazoa (cnidarian and bilaterian ancestor). While the apical domain GRN is extensively studied in NVE and compared with bilaterians, the evolutionary relationship of the apical domain within cnidarian groups remains unknown. Hear, we utilised the well-defined developmental GRN framework of NVE to reveal how these conserved regulatory interactions have shifted in the apical domain of different cnidarian groups. Despite the morphological diversity between Actiniaria and Scleractinia planula primarily lacking apical tuft, the AMI and ATE larvae share an extensive gene profile with NVE in the apical domain. The shared gene composition with NVE, thereby shared apical signalling system, reflects homology. Even though shared TFs unite Actiniaria and Scleractinia planula, the architecture of apical domain and ciliary transcriptome are considerably different.

Next, we showed that genes involved in patterning the apical domain of anthozoan larvae are mainly absent in the stem leading to Medusozoa planula, suggesting that the scyphozoan lacks apical organ homologues to anthozoans; this implies a dramatic reorganisation of GRN in Medusozoa apical domain. Along the anteroposterior axis, the AAU showed drastic changes in GRN specifying oral-aboral identity; several crucial apical organ and neuronal genes along the anteroposterior axis are lacking shared expression with NVE. Strikingly, most of these genes are evolutionarily conserved between anthozoans and bilaterians, associated with anteroposterior neurogenesis and axial formation. That suggests that AAU has undergone extended changes in GRN after the split from anthozoans. It might be early to determine this based on current transcriptome data from tissue-specific transcriptome alone. On the other hand, some of our results accord with previous findings in a hydrozoan species, *Clytia hemisphaerica*, suggesting that the Medusozoan larvae are indeed much simpler than the anthozoan.

The magnitude of variation between AAU and NVE makes it rather difficult to know how the common ancestor of the cnidarian planula looked in terms of morphology and apical domain organisation. However, our data provide an interesting perspective on the issue of the ancestral cnidarian planula before the Anthozoa–Medusozoa split. Previous studies in comparison of molecular data proposed that anthozoan polyps, medusozoan polyps and a jellyfish stage are equally different from one another and suggested that the only truly conserved stage among the Anthozoa and Medusozoa might be the planula larva, which becomes the best candidate for the cnidarian ancestral body plan (31). However, our data demonstrate that the planulae of anthozoans and medusozoans are also highly different.

With this study, we provide crucial insights into the molecular signature of the larval sensory structure across the Cnidaria and its evolutionary history. Based on the earlier studies in scleractinian and medusozoan planulae, it is clear that neuropeptide-expressing cells are present in all groups of cnidarians. We show that the scleractinian planulae share most apical domain GRN with NVE, suggesting that scleractinian planulae have apical cells homologous to NVE. On other hand, medusozoans potentially lack an apical organ that is homologous to anthozoans and bilaterians. It is plausible that the common ancestor of cnidarians possessed an apical plate comprised of apical and sensory-neurosecretory cells and that after the split of Actiniaria from other Anthozoans, the Actiniaria planula innovated cilia-associated genes leading to the origin of apical organ with a ciliary tuft **(Fig. 5E)**.

## Methods

### Animal collection and culturing

*Nematostella*: Polyps were grown in 16 ‰ artificial seawater at 18°C in the dark and fed with freshly hatched *Artemia* nauplii. The induction of spawning was performed as previously described (100). After fertilisation, the gelatinous substance around the eggs was removed using 4% L-Cysteine (Sigma-Aldrich, USA) (100). *Aurelia aurita*: Ephyras were collected using a plankton net in the vicinity of Plymouth Sound, UK and cultured in seawater at 18°C in a 12:12 light and dark cycle. The jellyfish were fed twice a day with freshly hatched *Artemia nauplii*. Fertilised embryos were collected from the brood sacs and reared to the planula stage. *Acropora millepora* and *Acropora tenuis*: The corals were cultured in ex-situ (101, 102). During the annual spawning of 2020, the embryos were collected after fertilisation. The larvae are maintained in artificial seawater at 27 °C and collected for experimentation at the planula stage.

### Microdissection of planula larvae

To separate the apical region from the rest of the larval body, we performed microdissection on *Nematostella, Aurelia aurita, Acropora millepora* and *Acropora tenuis* larvae as described in our previous study in *Nematostella* (33). Apical tissue containing the apical organ was isolated using 34-gauge needles under a stereomicroscope with 10X magnification. Motile larvae were placed into a fresh plastic Petri dish filled with *Nematostella* medium or filtered seawater. The larvae tend to adhere briefly to the bottom of a new plastic Petri dish, allowing enough time to separate the apical tissue by cutting. Each sample was pooled from a minimum of 100 individual larvae. For each planula prior to dissection, the frontal region is identified by means of the direction of larvae swimming. The samples were carefully collected using glass Pasteur pipettes, the excess medium was removed, and the samples were snap-frozen in liquid nitrogen and stored at -80°C until further processing.

### RNA sequencing and differential gene expression

For RNA isolation, due to the sheer size, samples collected from multiple batches were combined to acquire an adequate amount of RNA for sequencing. Total RNA was isolated using the TRI Reagent® according to the manufacturer’s protocol. RNA quality was assessed using Agilent RNA 6000 Nano Kit on Agilent 2100 Bioanalyzer (Agilent, USA), and samples with RNA integrity number ≥ 8.0 were used for sequencing. The CORALL RNA-Seq Library Prep Kit (Lexogen GmbH) was used for library preparation. Before sequencing, the libraries were pre-assessed by Agilent High Sensitivity DNA Kit (Agilent, USA) and quantified using Qubit™ 1X dsDNA HS Assay Kit (Invitrogen™). The sequencing was outsourced (GENEWIZ Illumina NovaSeq™ 2x150 bp sequencing), generating 15 million paired-end reads per replicate. Raw files and processed data were deposited at NCBI GEO submission GSE242174. After de-multiplexing and filtering high-quality sequencing reads, the adapter contamination was removed using *fastp* an ultra-fast all-in-one FASTQ preprocessor (103). Further, the quality of the reads was verified using FastQC (104). Processed reads from each sample were mapped to the respective genome and gene models (indexed bowtie2 (105)) by using STAR by using STAR (Spliced Transcripts Alignment to a Reference) (106). *Nematostella* gene model (https://figshare.com/articles/Nematostella_vectensis_transcriptome_and_gene_models_v2_0/807696). *Aurelia aurita* (107), Coral *Acropora millepora* (108) and *Acropora tenuis* (109). The number of reads mapping to the respective gene model were extracted from STAR output using the featureCounts tool (110). Differential expression analyses were performed using DESeq2 (Galaxy Version 2.11.40.7+galaxy2) (111).

### Identification of orthologs

We performed a BLASTP (112) search with the default curated gathering threshold for functional annotation to predict the protein homologs against the UniProt database (113). Additionally, we used gene functional annotation data from the published studies of respective species: *Nematostella* single-cell transcriptome study (61, 114) and *Aurelia* genome study (31, 98) **(Supplementary file 1)**. To identify genome-wide orthologous clusters across selected cnidarian species, we used OrthoVenn3 (https://orthovenn3.bioinfotoolkits.net/home) (115, 116), a genome-wide comparison and visualisation tool. Protein sequences of the reference genomes were organised into orthologous gene groups based on sequence similarity (116). The whole-genome protein sequences of all species were checked for sequences containing characters other than amino acids in the fasta file, and sequences smaller than <20 amino acids were removed. We retrieved the protein sequences of all gene coding sequences without alternative splice variants. The OrthoMCL performs an all-against-all BLASTP alignment, identifies putative orthology and in-paralogy relationships with the Inparanoid algorithm (117) and generates disjoint clusters of closely related proteins with the Markov Clustering Algorithm (MCL) (118). Further, to deduce the putative function of each ortholog, the first protein sequence in each cluster are subjected to BLASTP analysis against the non-redundant protein database in UniProt (113).

### Phylogeny

For FGF and PRD class homeobox phylogenetic analysis, the following species were selected from Actiniaria: *Nematostella vectensis, Exaptasia pallida*, Scleractinia: *Acropora millepora* and *Acropora tenuis*, and Medusozoa: *Clytia hemispherica* and *Aurelia aurita*. The protein sequences considered for the FGF and PRD class hox homologs search was provided in the supplementary file **(Supplementary file 1)**. We carried out BLASTP searches using the published transcriptome data of each selected cnidarian species. In case we found no matches, we carried out TBLASTN searches with the same query sequences. The sequences were obtained from orthology analysis or blast search. The protein sequences were aligned using MUSCLE (119) algorithm in the SeaView program (120). The maximum-likelihood (ML) phylogenetic trees were constructed using PhyML 3.0 online (www.atgc-montpellier.fr/phyml/) (121). The model was automatically selected by the Smart Model Selection with (SH-aLRT) (122). Statistical tree robustness was assessed in PhyML via 100 bootstrap replicates.

### In situ hybridisation (ISH)

ISH was performed according to published protocols (123, 124). In brief, fixed animals were transferred into sieves and rehydrated in 1 mL 60% methanol/40% PBST and then washed in 30% methanol/70% PBST. Samples were digested in proteinase K (80 µg/mL) for 5 min then blocked in glycine (4 mg/mL). Larvae were then transferred into 4% formaldehyde at RT for 1 h. Hybridisation was carried out with DIG-labelled probes for 48 h at 60 °C. After incubation, samples were washed through serial dilutions of 25%, 50%, 75%, 100% 2x SSCT at hybridisation temperature. The colour development was carried out in a 1:50 dilution of NBT/BCIP at RT. Stained animals were visualised with a Leica DM1000 microscope equipped with a MC190 HD Microscope Camera (Leica, Germany). For each gene at least 30 specimens were tested.

### Whole-mount immunofluorescence and SEM

After fixation, the samples were washed 5 times with PBST (1× PBS, 0.05% (vol/vol) Tween-20) for 10 min. The samples were blocked in 5% BSA in PBST for 1 hr at RT. Primary antibody (1:500 dilution, mouse Anti-α-Tubulin Cat # T9026, Sigma-Aldrich) incubation was performed in a blocking solution (1% BSA in PBST) for 24-36 hr at 4°C. The samples were washed with PBST for 5 x 5 min, after which samples were incubated with secondary antibodies (1:250 dilution; Goat anti-Mouse IgG Alexa Fluor 594 Cat # A-11032, ThermoFisher) diluted in blocking solution for overnight at 4°C. Then, the samples were washed with PBST for 5 x 10 min. Imaging was performed on Leica TCS SP8 DLS and Leica DMi8 confocal microscopes. Sample preparation for scanning electron microscopy was performed as in (125); SEM imaging was performed using the JEOL IT 300 scanning electron microscope.

## Acknowledgements

We thank Kevin Atkins for his help with setting up the sea anemone facility. We also thank Alix Harvey, Belle Heaton and Lucy Holloway for their support in maintaining the animal facility at the Marine Biological Association. We acknowledge Hartley George and Matthew Drysdale from Horniman Museum and Gardens for their input in rearing the coral larvae. We thank Glenn Harper and Dr Alex Strachan from Plymouth Electron Microscopy Lab for their assistance during the imaging.

## Author contributions

Conceptualisation: V.M.; Methodology: E.G., J.C., V.M.; Software: V.M.; Validation: E.G., V.M.; Investigation: E.G., V.M.; Resources: V.M.; Data curation: E.G., V.M.; Writing - original draft: V.M.; Writing - review & editing: E.G., Visualisation: E.G., V.M.; Supervision: V.M.; Funding acquisition: V.M.

